# Enhancing neuronal plasticity through intracranial theta burst stimulation in the human sensorimotor cortex

**DOI:** 10.1101/2021.03.29.437614

**Authors:** Jose L. Herrero, Alexander Smith, Akash Mishra, Noah Markowitz, Ashesh D. Mehta, Stephan Bickel

**Affiliations:** The Feinstein Institutes for Medical Research, Northwell Health, Manhasset, New York; Department of Neurosurgery, Hofstra Northwell School of Medicine, Manhasset, NY; Department of Neurology, Hofstra Northwell School of Medicine, Manhasset, NY

**Author notes:** **Correspondence:** Jose L. Herrero, Institute for Bioelectronic Medicine, The Feinstein Institutes for Medical Research, Northwell Health. 350 Community Drive, Manhasset, New York.

**Keywords:** Intracranial EEG, direct electrical brain stimulation, iTBS, neuronal plasticity, beta oscillations, sensorimotor cortex

## Abstract

The progress of therapeutic neuromodulation greatly depends on improving stimulation parameters to most efficiently induce neuroplasticity effects. Intermittent Theta Burst stimulation (iTBS), a form of electrical stimulation that mimics the natural brain activity patterns, has shown efficacy in inducing such effects in animal studies and rhythmic Transcranial Magnetic Stimulation (rTMS) studies in humans. However, little is known about the potential neuroplasticity effects of iTBS applied through intracranial electrodes in humans which could have implications for deep brain stimulation therapies. This study characterizes the physiological effects of cortical iTBS in the human cortex and compare them with single pulse alpha stimulation, another frequently used paradigm in rTMS research. We applied these stimulation paradigms to well-defined regions in the sensorimotor cortex which elicited contralateral hand or arm muscle contractions during electrical stimulation mapping in epilepsy patients implanted with intracranial electrodes. Treatment effects were evaluated using effective connectivity and beta oscillations coherence measures in areas connected to the treatment site as defined with cortico-cortical evoked potentials. Our results show that iTBS increases beta band synchronization within the sensorimotor network indicating a potential neuroplasticity effect. The effect is specific to the sensorimotor system, the beta frequency band and the stimulation pattern (no effect was found with single-pulse alpha stimulation). The effects outlasted the stimulation by three minutes. By characterizing the neurophysiological effects of iTBS within well-defined cortical networks, we hope to provide an electrophysiological framework that allows clinicians and researchers to optimize brain stimulation protocols which may have translational value.

## Introduction

Brain stimulation therapies such as transcranial magnetic stimulation (TMS) and electrical deep brain stimulation (DBS) of subcortical and cortical structures are increasingly used to treat neurological and psychiatric disorders such movement disorders [1], epilepsy [2,3], major depressive disorder [4,5], OCD [6], and Tourette’s syndrome [7], and they are actively being studied for use in stroke recovery [8], PTSD [9], and substance abuse disorders [10,11]. In DBS, the therapeutic effect is thought to be related to acutely enhancing or inhibiting activity in specific brain regions and until recently, less emphasis was placed on the potential contributing role of neural plasticity induced by electrical stimulation. Despite the important role of neural plasticity as part of the therapeutic effects in DBS [12–14], there is still much to learn about the most efficient plasticity inducing stimulation parameters using this approach [15].

Non-invasive stimulation modalities such as repetitive TMS can cause systematic changes in cortical excitability [16]. For example, high-frequency rTMS (> 5 Hz) applied to the motor cortex caused increased cortical excitability as measured by the amplitude of motor-evoked potentials (MEP) in contrast to low-frequency (~1 Hz) stimulation which more frequently led to an opposite effect [16–18]. While rTMS is currently used to treat a wide range of clinical conditions, TMS-induced plasticity effects are transient, require repeated treatment visits and are effective in approximately 30% of patients [18,19]. Thus, the portability and addition of potential anatomical targets renders DBS as a valid treatment option in certain patient populations despite its invasiveness [2,20].

A recent trial that tested high-frequency rTMS-like stimulation applied directly to the brains of individuals with intracranially implanted electrodes, found neuroplasticity effects in functionally connected brain areas [21]. However, contrary to the rTMS findings, this trial demonstrated subject-dependent and site-dependent responses (enhancement or suppression) that could not be predicted by the characteristics of the stimulation frequency alone. A potential explanation of the heterogenous effects is that stimulation was applied across several different functional regions across patients. Additionally, the authors used intermittent alpha burst stimulation while the most efficient protocol to induce increases in neural excitability uses intermittent theta burst stimulation (iTBS, brief bursts of 50-100Hz pulses repeated at 5 Hz) [18,22–28]).

In our study we directly applied iTBS to one specific network, the sensorimotor, and assessed the treatment effects exclusively on connected sites as defined by cortico-cortical evoked potentials (CCEPs) [29–33]. The treatment sites were carefully selected based on clinical functional stimulation mapping results in seven patients that had extensive sensorimotor cortical coverage. As a read-out of the effect, we measured changes in effective connectivity using CCEPs (similar to [21]) as well as changes in beta coherence across the sensorimotor network during the periods preceding and succeeding the treatments. Sites within the sensorimotor network exhibited prominent beta oscillations, motivating our coherence analyses within the beta frequency range. These results might guide further investigations in the design of stimulation protocols aiming at inducing neuroplasticity in cortical networks.

## Methods

### Patient selection

Seven patients with medically intractable epilepsy who underwent electrodes implantation at North Shore University (six) or Lenox Hill (one) hospitals for seizure localization were enrolled in the study. The decision to implant, the electrode targets, and the duration of implantation were made entirely on clinical grounds. The patients had sensorimotor electrode coverage because the clinical hypothesis included a potential involvement of these areas in the seizure network. Patient characteristics are described in Table 1. Patients were selected on the bases of having 1) ample electrode coverage of sensorimotor areas; 2) contralateral hand and/or arm motor contractions upon clinical high frequency stimulation mapping (HFSM) and 3) confirmed seizure onset focus outside sensorimotor areas. All patients provided informed consent as monitored by the local Institutional Review Board following the Declaration of Helsinki. Patients were informed that participation in this study would not alter their clinical care, and that they could withdraw from the study without jeopardizing their care. Three patients in this study were included in a prior study [34]. However, the evaluations put forth here are unique to this study as they refer to different stimulation sessions and stimulation parameters.

**Table 1.**
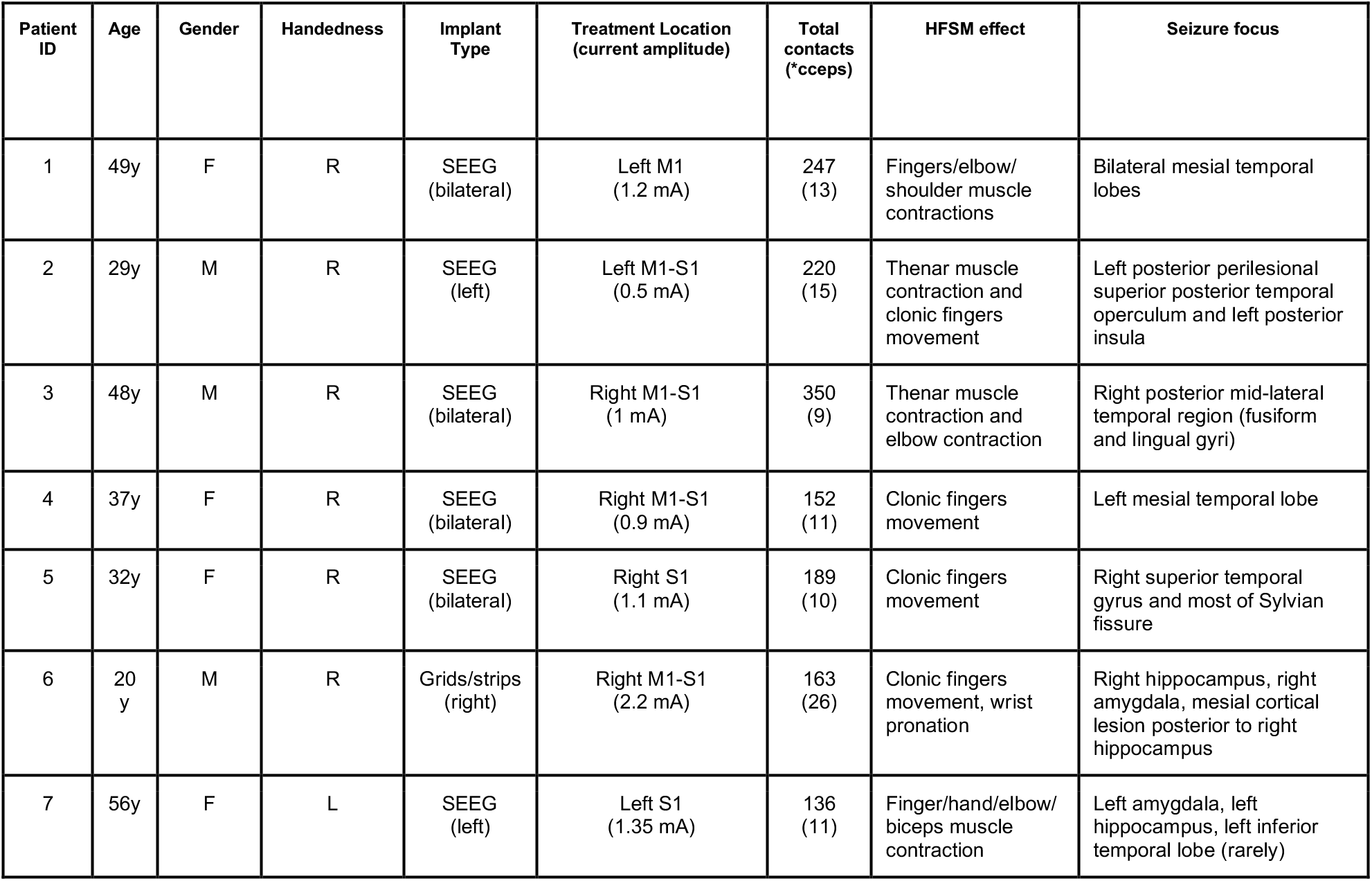
Summary of patients and stimulation characteristics.

**Table 1.** High Frequency Stimulation Mapping (HFSM): clinical procedure during which every electrode contact pair is stimulated with high frequency stimulation bursts to determine function. All HFSM-related motor contractions were contralateral to the stimulation site. Treatment locations for iTBS/i8Hz were selected based on HFSM results (contact pairs that elicited the most specific finger/arm contractions) and the final current for the subsequent treatment was set at an intensity of 80% of the active motor threshold. Treatment locations: M1, primary motor cortex (both stimulation contacts were immediately anterior to the central sulcus); S1 primary somatosensory cortex (both stimulation contacts were immediately posterior to the central sulcus); M1-S1 Sensorimotor (paracentral, one stimulation contact was anterior to the central sulcus and the other posterior to it). M-Male; R-Right; L-Left; SEEG-stereo EEG. Total contacts: total number of electrode contacts recorded and total number of contacts with significant CCEPs located in grey matter (white matter and cerebrospinal fluid contacts were discarded). Seizure Focus-all seizure onset foci were outside motor/premotor areas.

### Electrode registration

Our electrode registration method was described in detail previously [35]. Briefly, we utilized the iELVis toolbox, which makes use of BioImage Suite, FSL, FreeSurfer, and custom written code for intracranial electrode localization. The electrode contacts are semi-manually located in the postimplantation CT which is co-registered to the preimplantation MRI. Additionally, FreeSurfer aligns the patient’s pre-implantation MRI to a standard coordinate space, automatically parcellating the brain and assigning anatomical regions to each contact. Finally, iELVis software is used to project the contacts locations onto FreeSurfer’s standard image.

### Electrophysiological recording and preprocessing

iEEG signals were acquired continuously at 512 Hz or 3 kHz using a clinical recording system (Xltek Emu 256, Natus Medical) or a Tucker-Davis PZ5M module (Tucker Davis Technologies). iEEG data was extracted by bandpass filtering (8-pole Butterworth filter, cutoffs at 0.01 and 250 Hz). Either subdural or skull electrode contacts were used as references, depending on recording quality at the bedside, and were subsequently re-referenced to a common average. The power spectrum of the signals was inspected online before the experiment started to ensure its physiological properties.

### High frequency stimulation mapping (HFSM)

HFSM is a clinical functional mapping technique to localize seizure onset (and eloquent) areas which is performed in intracranial patients after anticonvulsant medications are resumed. A Grass S12D (Grass Technologies) or a Tucker-Davis (IZ2MH) stimulator was used to apply bipolar stimulation with biphasic matched-square wave pulses (100 μs or 200 μs/phase), and current amplitudes ranging from 0.5-4 mA or 0.5-10 mA for depth and subdural electrodes respectively, at a rate of 50 Hz up to 0.5-2 seconds duration. All seven patients included in this study had HFSM of sensorimotor cortical sites which elicited contralateral clonic or tonic-clonic muscle contractions (Table 1).

### Treatment site selection and stimulation protocol

As treatment sites we selected electrode contacts in which HFSM elicited the most specific contralateral finger or hand movements (Table 1) at the lowest current amplitude (e.g., active motor threshold). To minimize variability across patients the preferred stimulation site was the hand area of the motor cortex. However, sites of electrode implantation were based upon clinical criteria, and not every patient had electrodes implanted within the hand area. In these cases, the closest region to the hand motor cortex that elicited finger, wrist or arm movements was selected. Treatment stimulation currents matching 80% of the active motor threshold were used as in previous studies [17,24,25], and no epileptiform after-discharges occurred at these intensities. iTBS in TMS studies consists of three pulses delivered at 50 Hz (20-ms separating each pulse), and each set of three pulses is repeated at 5 Hz (Fig. 1, inset). In our iTBS protocol we applied this sequence of pulses for two seconds (30 pulses/train) repeated every ten seconds (8-s intratrain interval). Each pulse consisted of a bipolar, bi-phasic, square-wave with 200 μs/phase. In the comparison condition (i8Hz), single stimulation pulses were administered at intervals of 125-ms for a period of five seconds (40 pulses/train) and repeated every fifteen seconds (10-s intratrain interval). Similar stimulation frequencies ranging from 3-8 Hz were used in recent DBS studies [15,36,37]. In our study, the number of total pulses was similar across treatment modalities (iTBS, 30*20=600 pulses; i8Hz, 40*15=600 pulses). Treatments were applied in a randomized order across subjects with a long (>45 minutes) interval between treatments to allow for the washout of any residual effects from the previous treatment. Resting state activity immediately before and after each treatment was recorded to compute the coherence measures (Fig. 1, Inset).

**Fig. 1.**
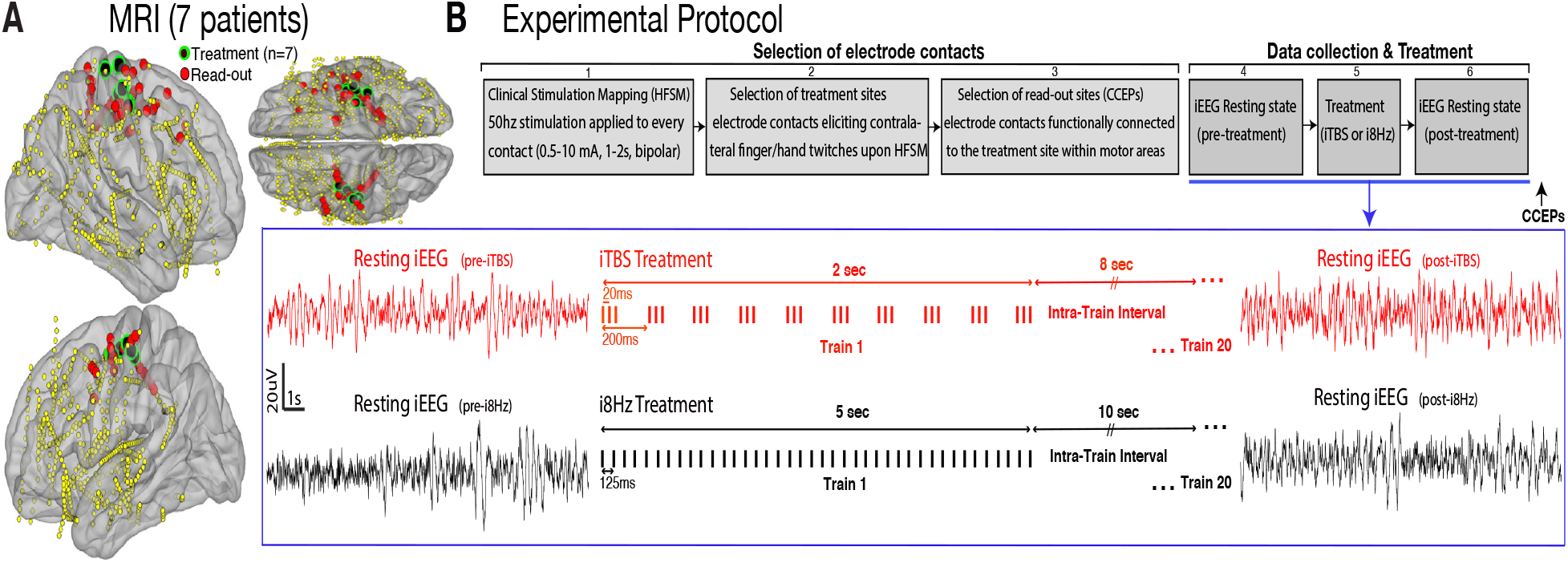
Treatment locations and experimental protocol. **(A)** Electrode contacts from 7 subjects registered onto a common brain surface. Treatment locations (green dots). Read-out electrode contacts (red dots) are connected to the treatment locations based on CCEP mapping. Only sensorimotor contacts are shown here. **(B)** Experimental protocol. Box 1, HFSM reveals sites in sensorimotor cortex that elicit specific contralateral motor responses upon stimulation. Box 2, calculation of motor thresholds for each patient and sites of interest. A stimulated site eliciting the lowest threshold for motor response was chosen as the treatment site. Box 3, application of single pulses to treatment sites (>150 pulses, 4mA, 200μsec pulse width) and selection of read-out sites (contacts connected to the treatment site) based on CCEP analyses (see Fig. 2a). Box 4, recording of iEEG resting activity while subjects are awake but quiet and avoiding movements (2-minutes). Box 5, treatment is applied (either iTBS or i8Hz, see inset). Box 6, recording of iEEG resting activity immediately after treatment ends (2-minutes). Single pulses to treatment sites (~200 CCEPs) were also administered after the post-treatment resting period (black arrow) to assess potential treatment effects on CCEPs amplitude. **Inset:** Top) ITBS treatment: Three pulses of stimulation administered at 50 Hz repeat every 200ms for a period of 2s and an 8s intra-train interval (2s ON and 8s OFF). Each 2s iTBS train contains 30 pulses and repeats 20 times (30*20 = 600 pulses total). Bottom) i8Hz treatment: stimulation pulses administered at 8 Hz (one pulse every 125ms) for a period of 5 seconds (40 pulses per train) and a 10 seconds intra-train interval (5 sec ON and 10 sec OFF) and repeats 15 times (40*15 = 600 pulses total).

### Cortico-cortical evoked potentials (CCEPs)

To localize brain areas connected to the treatment sites we performed CCEP mapping (Fig. 2a) [30]. CCEP mapping has been used to examine fronto-parietal [29,38], hippocampal [39], visual [40], language [31,32,41] and other networks [38]. Single pulse electrical stimulation (biphasic square-wave pulses (200 μs/phase) were applied at the treatment site (200 pulses, 1s interstimulation interval with a +/-300ms jitter). To get robust CCEPs, stimulation currents were set at 4 mA or just below the threshold that elicited movement. We also used CCEPs to probe for possible neuroplasticity effects and applied CCEP stimulation before and after treatment (iTBS or i8Hz, Fig. 1b). CCEP stimulation ended at least 15-minutes prior to treatment onset to avoid potential residual effects on neuronal excitability. Statistical significance of CCEPs was determined as follows: 1) a bipolar montage (first spatial derivative) was applied to the data to reduce volume conduction effects; 2) data from each electrode contact were epoched −1000 to 1000 ms centered on the electrical pulse; 3) single CCEP traces were demeaned, baseline corrected (−200 to −25 ms) and averaged; 4) the averaged CCEP trace was transformed to a t-statistic by dividing the average at each time point by the standard error of the mean at the same time point. The absolute maximum of the t-statistic during the post-stimulus period (10-50 ms) was compared against the distribution of the t-statistic during the pre-stimulus baseline period. If the post-stimulus t-statistic was higher than 6 standard deviations, the response was considered as significant. These thresholds (6 SD) and time windows (10-50 ms, when the most direct synaptic effects occur) were based on prior studies [33,39]. Significant electrode contacts were considered as read-out contacts in which treatment effects were later evaluated. Read-out contacts outside gray matter (in white matter and cerebrospinal fluid) were discarded from the main analyses of treatment effects.

**Fig. 2.**
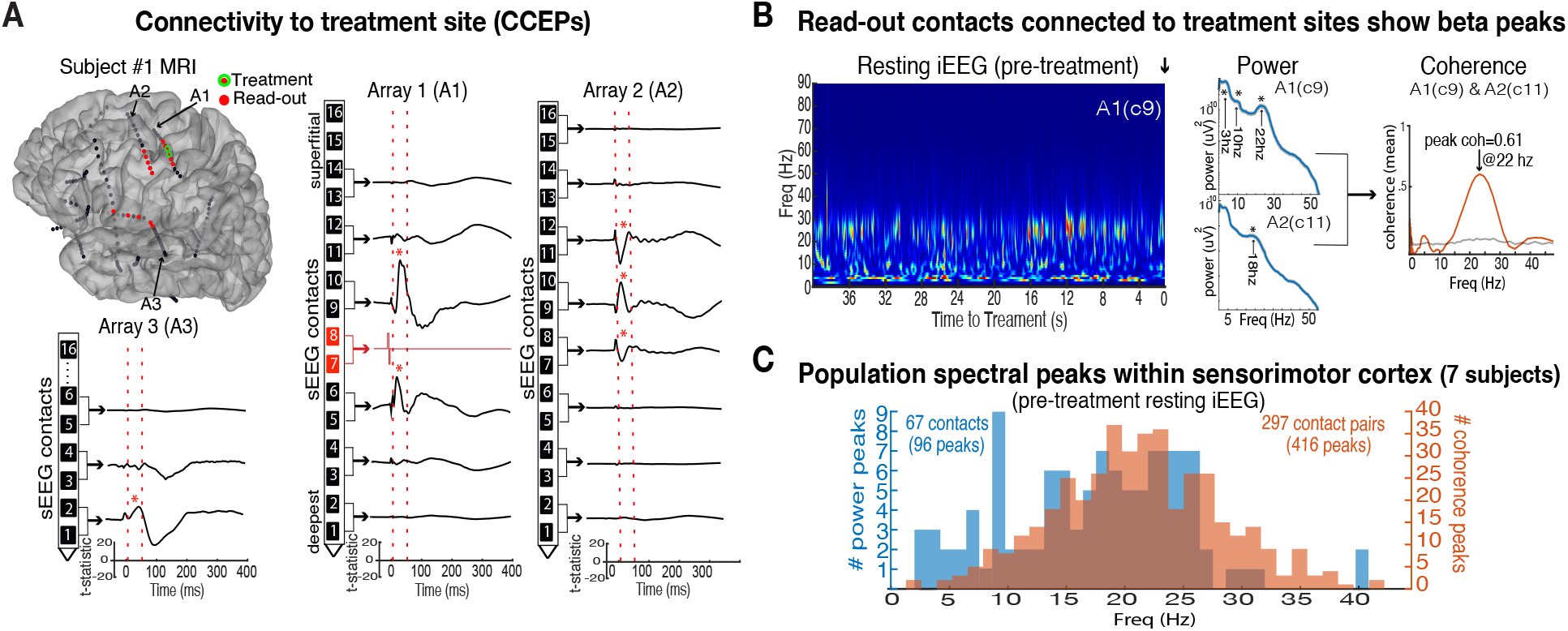
Areas connected to treatment sites have high-beta oscillations. **(A)** Connectivity to treatment sites using cortico-cortical evoked potentials (CCEPs) in subject 1; single pulses (200 sweeps, bipolar stimulation, 4mA) applied to the treatment site (Array 1, contacts 7-8, red line) elicit strong evoked potentials at nearby sites within sensorimotor cortex (A1 contacts 5-10 and A2 contacts 7-12) and at more distant sites (A3 with contacts 1-4 located in the insula, and other more distant contacts located in the most lateral portion of the sensorimotor cortex). Asterisks marked contacts with significant CCEPs (P1 peak amplitude > 6SD t-statistic). Vertical dashed lines (red); analyses window (10-50ms post-stimulation). **(B)** (Left) Time-frequency power spectrum of one example electrode contact located in the motor cortex adjacent to the treatment site (A1, c9) during a 2-minutes resting period recorded prior to treatment. (Center) Averaged spectrograms show prominent spectral peaks in the beta band for the same contact (top, ~22 Hz peak) and another contact adjacent to it (bottom, 18 Hz peak). (Right) Coherence between these two contacts (orange line, mean coherence across all overlapping windows; grey line pointwise significance level, ***α*** = 0.05). **(C)** Population spectral peaks (67 electrode contacts) and coherence peaks (297 contact pairs) of connected sites based on CCEP mapping within the sensorimotor cortex across all patients; power peaks (blue bars, 96 peaks from 67 contacts) and coherence peaks (orange bars, 416 peaks from 297 contacts).

### Coherence and power analyses

Analyses were conducted using the FieldTrip toolbox [42] and custom MATLAB scripts. After determining the sites of interest using CCEPs (e.g., contacts functionally connected to the treatment site, Fig. 2a), we applied a sensor-based analysis to these contacts. Given the caution against the use of bipolar EEG for synchronization analyses [43–45], we used common average reference montage for our coherence analyses. To calculate the coherence between two iEEG contacts, we squared the magnitude of the cross-spectrum of their raw signals, normalized it by the power spectra of each signal at each respective frequency, and smoothed it [46]. The imaginary part of the coherence was used for the analysis as it might be less impacted by common input sources [47]. Each 2-min data set (e.g., pre-treatment) was divided into overlapping 10-s windows (1/10 = 0.1-Hz resolution, sliding 2-s at a time) and windows with outliers (> 4 SD of the mean coherence across all sliding windows) were discarded. The total windows discarded across all subjects were 0.8%. Statistical significance of dominant spectral coherence peaks was evaluated using cluster-based non-parametric tests implemented in Fieldtrip (see Fig. 2c population coherence peaks). iEEG signals between contact pairs were shuffled (1000 iterations) and the critical levels of the Wilcoxon-Mann-Whitney tests at each frequency were computed (see Fig. 2b, coherence plot for an example site, grey line shows pointwise significance level, ***α*** = 0.05, jackknife method). The dominant peaks in the power spectrum were determined using power spectral density estimates using the “periodogram.m” function (MATLAB) with 95% confidence bounds (Fig. 2b, example power plots; Fig. 2c, population power peaks). To assess whether the coherence between iTBS/i8Hz pre-treatment and post-treatment was significantly different for any given contact pair (and at what specific frequencies), we used the non-parametric Monte Carlo test implemented in “ft_freqstatistics.m” at alpha-level 0.01 (Fig. 3a, red line above x-axis). The peak coherence values at the beta band (12-30 Hz) for all sensorimotor contact pairs combinations were extracted and compared in each patient (pre vs. post). We used paired nonparametric tests to evaluate differences between matched samples in iTBS and i8Hz (Wilcoxon signed rank tests, p < 0.01) (Fig. 3b, 4). Repeated measures ANOVA were used to assess the effect of the interaction (treatment (iTBS vs. i8Hz) * time (pre vs. post), p < 0.05). Similar statistical tests were used to compare the spectral power across treatments. The coherence in the intra-train analyses (Fig. 5d) was calculated similar as the coherence during the pre/post-treatment periods except that the analyses window was 6 seconds duration (+0.1 to 6.1-s after each 2-s train offset), sliding 500-ms at a time (1/6 = 0.16-Hz resolution). The peak coherence in the beta band (12-30 Hz) was determined after each train (#1-20) and a linear regression was fitted to the resultant values to compute the slope of the function. Statistical difference between slopes across treatment conditions was evaluated using paired nonparametric tests (Wilcoxon signed rank tests, p < 0.01) (Fig. 5e).

**Fig. 3.**
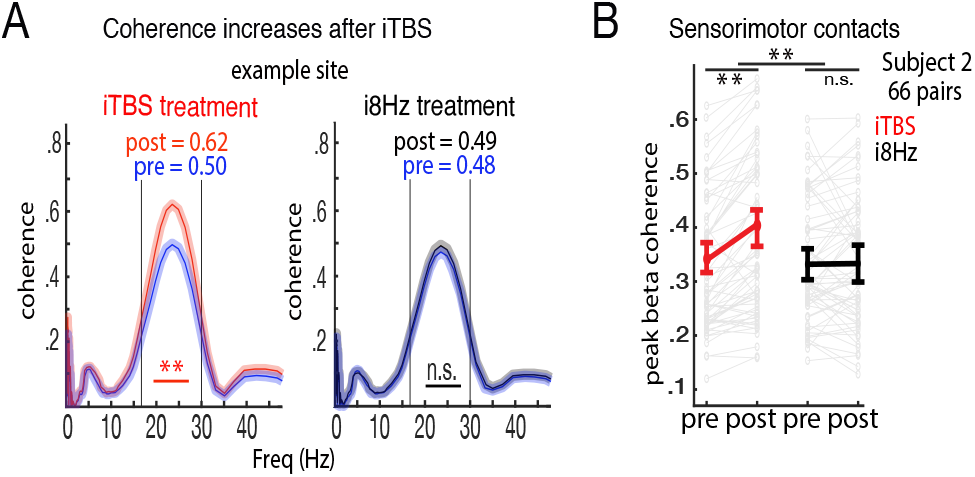
iTBS increases beta coherence in sensorimotor cortex while i8Hz does not. **(A)** Treatment effects for an example site located in the motor cortex of patient 2. Beta coherence is enhanced after iTBS but not after i8Hz (red line pointwise significance level marked with asterisks, *α* = 0.01; shaded areas indicate SE). **(B)** Averaged peak beta coherence for all sensorimotor pair combinations before and after treatments (iTBS, red line; i8Hz, black line) in patient 2 (66 pairs). Grey shade lines show the mean coherence peak for each single pair of contacts.

**Fig. 4.**
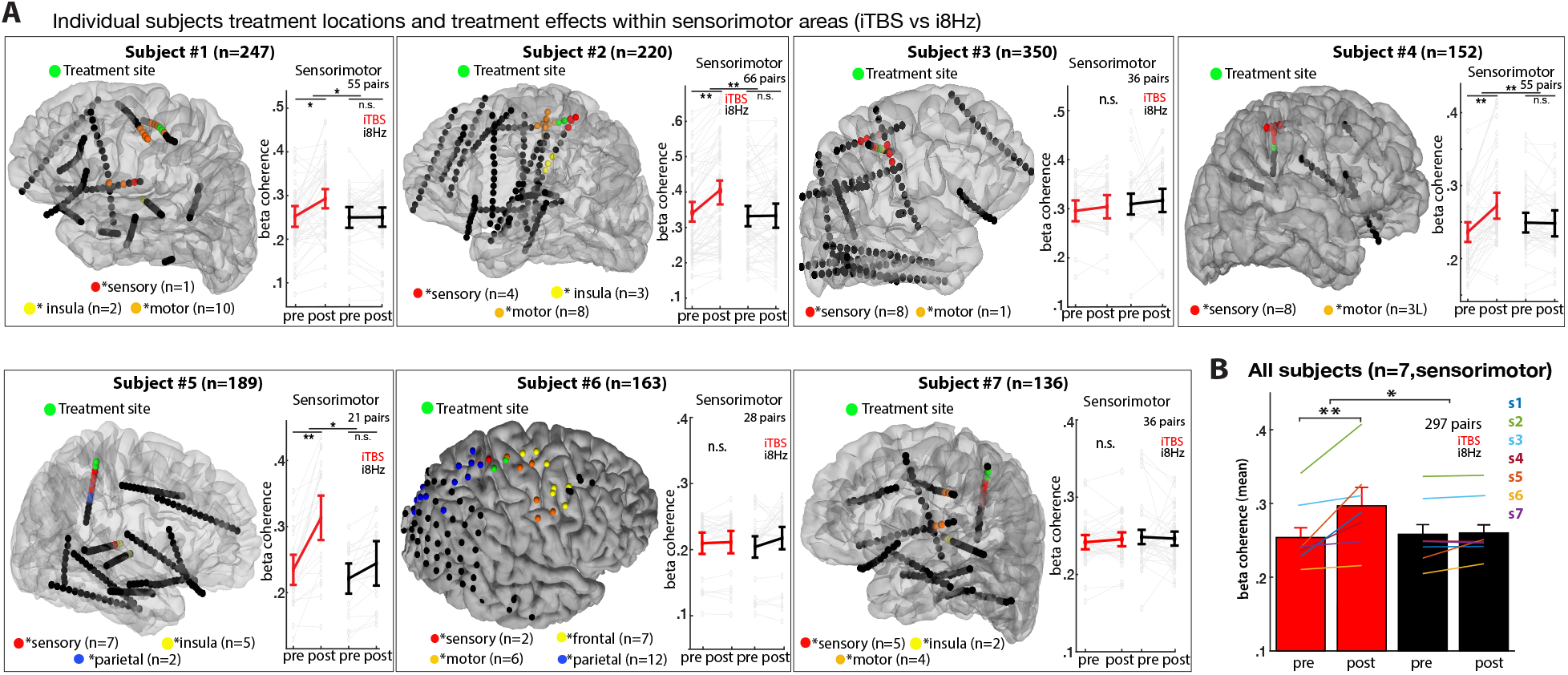
iTBS increases beta coherence in the sensorimotor cortex across the population data. **(A)** Individual subjects MRI and electrode locations. Treatment sites (green dots). Read-out sites (colored dots) represent electrode contacts with significant CCEPs when stimulating the treatment site (contacts with effective connectivity to the treatment site). Most read-out contacts were close to the treatment site in the somatosensory (red) and motor (orange) cortices, but also more distally in the insula (patients 1, 2, 5, and 7, yellow dots), parietal (patients 5 and 6, blue dots) and frontal cortices (patient 6, yellow dots). **Inset plots**: Individual subjects beta coherence for all contact pairs in the sensorimotor cortex before and after treatments (thin gray lines show the coherence for each contact pair and thicker lines, the averaged coherence for iTBS (red) and i8Hz (black). iTBS increases beta coherence in the sensorimotor cortex in 4/7 subjects. Significant effects are marked with asterisks. **(B)** Population data: averaged beta coherence for all sensorimotor electrode contacts pairs across the 7 patients (297 pairs). Overlaid colored lines show the averaged beta coherence for each patient individually. Asterisks mark significance effects (* p < 0.05; ** p < 0.01; n.s., non-significant).

**Fig. 5.**
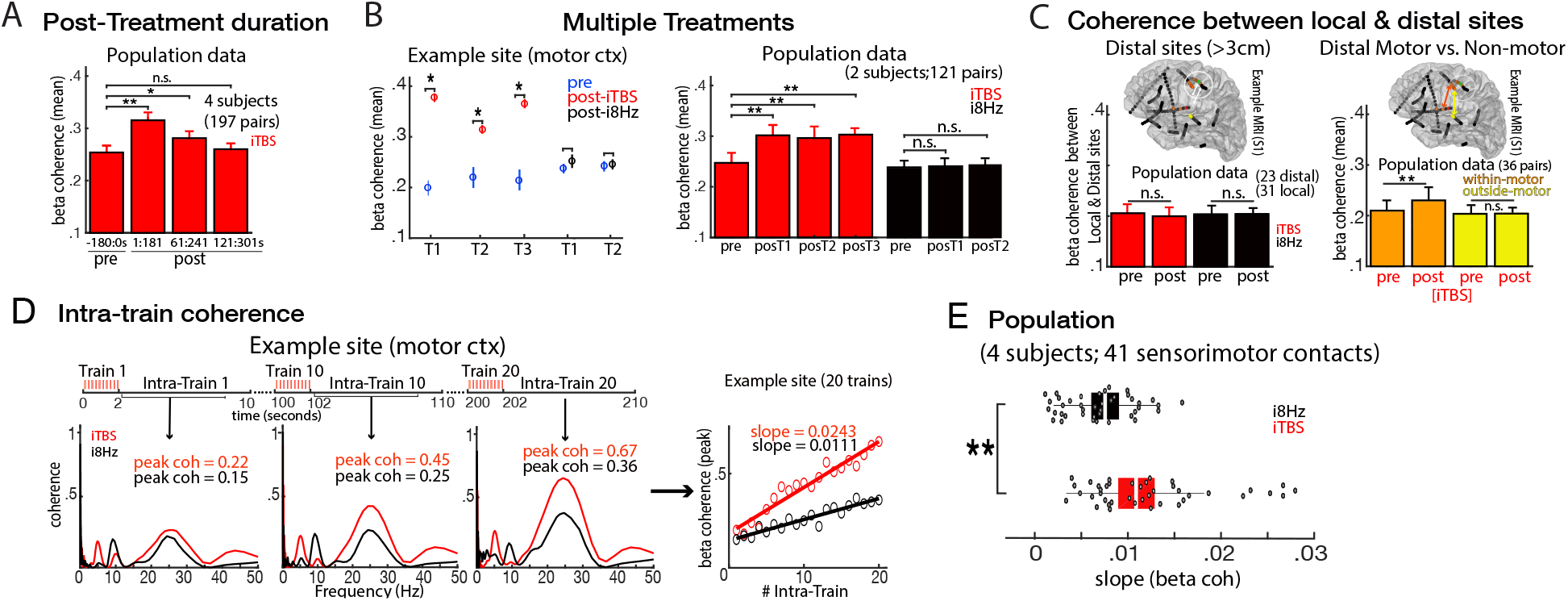
iTBS effects are frequency, spatially and temporally specific. **(A)** Post-treatment duration after-effects: iTBS-induced increases in beta coherence outlasted the duration of stimulation by three minutes -strongest effects occur during the initial 2 minutes (0-2 min), mild effects during 1-3 min and washed 2-4 min after treatment offset. Significant effects are marked with asterisks. **(B)** Multiple treatments effects; (Left) sequential application of iTBS treatments up to three times (3*600 total pulses, 5 minutes gaps in-between) does not further increase beta coherence for an example contact pair in the motor cortex (iTBS, red dots; pretreatment baseline, blue dots; i8Hz, black dots). (Right) Averaged data from all sensorimotor contact pairs in two patients (n = 121). **(C)** Coherence between local and distal sites: (Left) Example MRI brain from patient 1; white circle shows the 3 cm boundary around the treatment site. Contacts outside this boundary were considered distal (e.g., contacts in the insular, parietal, and most lateral and contralateral sensorimotor cortices). Bar plot shows the averaged beta coherence between local and distal contacts (4 patients) which was not affected by either treatment. (Right) MRI image depicts the coherence between focal and distal contacts that are still within sensorimotor cortices (orange arrow) vs. those distal contacts that are outside the sensorimotor cortex (yellow arrow). Bar plot shows the averaged beta coherence for the population data (iTBS treatment only) between local and I) distal contacts within sensorimotor cortex (orange bars) and II) distal contacts outside sensorimotor cortex (yellow bars). Coherence within sensorimotor areas was larger (9 contacts within vs. 9 contacts outside the sensorimotor area; 36 contact pairs; note the equal length of the orange and yellow arrows showing that the distance between pairs was matched across groups). **(D)** Intra-train effects: Buildup of coherence within a stimulation block for iTBS and i8Hz. (Left). Example contact pair within the motor cortex showing increases in beta coherence peak values from trains 1, 10 and 20. (Right) Linear function fitted to the beta coherence peaks from trains 1-20 shows coherence increasing linearly with steeper slope in iTBS. **(E)** Slopes across the population data (subjects 1, 2, 4 and 5, sensorimotor contacts, n = 41) are larger in iTBS compared to i8Hz. T1 = Treatment 1; pre = pre-treatment; posT1 = post-treatment 1.

## Results

### Treatment Sites

All seven patients selected for this study had ample electrode coverage of cortical sensorimotor areas (Fig. 1a & 4a). Selection of treatment sites was based on their function. For each patient, the electrode contact pair that elicited the most selective motor responses during HFSM was selected as a treatment site. Isolated (thumb and/or first) finger movements contralateral to the site of stimulation were elicited in 5 patients while additional recruitment of wrist and biceps was elicited in 2 patients (Table 1). Contact pairs were in the primary motor cortex (both stimulating contacts anterior to the central sulcus, patient 1), in the primary somatosensory cortex (both contacts paracentral, immediately posterior to the central sulcus, patients 5 and 7), and in the sensorimotor cortex (one contact immediately posterior to the central sulcus and the other anterior to it, patients 2, 3, 4 and 6) (Fig. 1a, 4a). Seizure onset foci was confirmed outside the sensorimotor areas in all patients (Table 1).

### Read-out sites

After determining the treatment site in each patient, we chose read-out electrode contacts based on significant responses during CCEP mapping. From a total of 1250 contacts recorded across 7 patients, 67 passed our selection criteria of exceeding a t-statistic > 6SD 10-50 ms post stimulation (35 in sensory/paracentral, 28 in motor and 4 in premotor cortices, Fig. 1a and 4a). An additional 26 contacts also exhibited significant CCEPs but were outside the sensorimotor regions. Figure 2a shows the results for an example subject. Single pulse stimulation applied to the treatment site (electrode array A1, contacts 7-8, located in the medial motor cortex) elicited significant CCEPs in contacts near to the stimulation site (A1 contacts 5-10, and A2 contacts 7-12), but also in more distant contacts (A3 with the deepest contacts located in the insula, and another array with contacts located in the most lateral aspect of the sensorimotor cortex). One of our outcome measures was treatment related changes in beta coherence because beta oscillations are the dominant spontaneously occurring network activity in the sensorimotor system. Figure 2 illustrates that sensorimotor contacts activities in our patients exhibited prominent beta peaks during resting-state prior to treatment. The averaged power spectra for two example contacts that were functionally connected to the treatment site within motor areas is shown in Fig. 2b. These contacts exhibited prominent spectral and coherence peaks at the beta frequency (12-30 Hz). The dominant spectral power peaks and coherence peaks for the population of sensorimotor sites across the 7 patients is shown in figure 2c. Spectral frequency peaks in power and coherence at other frequencies were also present in some sites together with the beta peaks.

### Treatment Effects of iTBS vs. i8Hz stimulation

Figure 3a shows an example electrode contact pair from the 12 read-out sensorimotor contacts selected (based on CCEP mapping) in one example subject. This contact pair shows increased beta coherence after iTBS treatment compared to before treatment (Wilkinson signed-rank test, p < 0.01). Beta coherence after i8Hz did not significantly change (Wilkinson signed-rank, p = 0.29). Figure 3b shows the coherence results across all contacts pairs combinations in this patient (n = 66) showing an increased coherence in the beta band after iTBS compared to before (Wilkinson signed-rank, p < 0.001) and a significant interaction between treatment (iTBS vs. i8Hz) and time (pre vs. post-treatment) (RM-ANOVA, F = 6.1, p = 0.0012).

Figure 4 shows the results across the population data for the 7 patients. From a total of 93 contacts with significant connectivity (based on CCEP mapping), we selected the 67 located in the sensorimotor cortex (39 contacts posterior to the central sulcus and 28 anterior to it) for our analyses of coherence (297 contact pairs combinations total). iTBS increased beta coherence in the sensorimotor cortex in 4/7 patients while i8Hz did not increase beta coherence in any of the patients. A repeated measures ANOVA with treatment (iTBS vs. i8Hz) and time (pre vs. posttreatment) across contacts (67 contacts with significant CCEPs located within the sensorimotor cortex across all 7 patients) showed a significant interaction (F = 6.34, p = 0.025). This result was driven by an increased coherence in the beta band after iTBS compared to before (Wilkinson signed-rank, p = 0.002). Individually this effect was significant for patients 1, 2, 4 and 5 while for patients 3, 6 and 7, the effect did not reach significance (Wilkinson signed-rank, both p > 0.1).

Beta coherence before and after the i8Hz treatment did not differ significantly at the population level (Wilkinson signed-rank, p > 0.1, all patients), nor for any of the patients individually.

### Treatment effect is frequency specific

The iTBS-induced effects were specific to the beta frequency band. Across all subjects no treatment effect was found in the frequency bands alpha (8-12 Hz), gamma (70-150 Hz) and theta (4-7 Hz) (both p > 0.4; data not shown). However, varying effects were found in the theta and alpha bands in individual patients. For example, iTBS treatment increased theta coherence in two patients (Wilkinson signed-rank, both p < 0.05), it reduced alpha coherence in one patient (Wilkinson signed-rank, p = 0.003), while no significant changes were observed in the other four (both theta and alpha bands, p > 0.4). The i8Hz treatment increased coherence in the alpha band (8-12 Hz) in one patient (Wilkinson signed-rank, p = 0.002), while no significant changes were observed in the other six (Wilkinson signed-rank, both p > 0.2).

### Treatment effect lasts approximately 3 minutes

The increase in beta coherence was strongest immediately after the entire block of iTBS treatment (+1:181 sec post-treatment compared to −180:0 sec pretreatment, Wilkinson signed-rank, p < 0.001) (Fig. 5a). The increased coherence effect was still significant but weaker during the 1-3 minutes post-treatment period (+61:241 sec, pre vs. post, Wilkinson signed-rank, p = 0.015), and it washed out for the 2-4 minutes post-treatment period (+121:301 sec, pre vs. post, Wilkinson signed-rank, p = 0.18).

### Multiple treatments do not result in further coherence increments

Next, we tested in two subjects whether repeated application of iTBS treatments resulted in gradual increments in beta coherence. Figure 5b shows the effects of 3 consecutive iTBS treatments (3*600 pulses separated by a 5-minute inter-treatment interval) for a pair of contacts in the motor cortex (left) and across the population data (right, 2 subjects, 121 contact pairs within sensorimotor networks). The increase in beta coherence after the first iTBS treatment was very similar compared to the second and third treatments (pre vs. post, Wilkinson signed-rank, both p < 0.001). Multiple i8Hz treatments did not significantly change beta coherence (pre vs. post, Wilkinson signed-rank, both p > 0.2).

### Treatment effect is specific to sensorimotor areas

We quantified whether the enhanced beta coherence after iTBS treatment observed in patients 1, 2, 4 and 5 was limited to read-out contacts near the treatment site (e.g., connected contacts within sensorimotor cortices < 3 cm from the treatment site) or extended to other, more distant yet connected (based on CCEP mapping) regions such as the insula (Fig. 5c, left). Distal contacts were in the insular (10), parietal (2) and more lateral aspects of the motor (8) and somatosensory (3) cortices and were averaged across areas and subjects before statistical testing (all 4 subjects had distal contacts including #4 in the opposite hemisphere; Fig. 4a). Beta coherence between focal (n = 31) and more distant contacts (n = 23) was not significantly changed after either treatment (pre vs. post, Wilkinson signed-rank, both treatments p > 0.1; RM ANOVA, stimulation*time interaction, F = 0.6, p = 0.41). These results suggest that iTBS-enhancement effects are relatively local within sensorimotor regions (< 3 cm from the treatment site). To examine the focal vs. network specificity of the iTBS-enhancement effect, we selected contacts within sensorimotor areas and compared them to other equally distant contacts that were outside sensorimotor areas. Contacts within sensorimotor areas located in the more lateral aspect of the sensorimotor cortices (> 3 cm away from the treatment site, 9/67) showed enhanced beta coherence compared to other equally distant contacts outside sensorimotor areas (Fig. 5c, right; distal sensorimotor contacts, pre-iTBS vs. post-iTBS, Wilkinson signed-rank, p = 0.019; distal non-sensorimotor contacts, pre-iTBS vs. post-iTBS, Wilkinson signed-rank, p = 0.36). This result suggests a network specific iTBS-enhancement effect rather than just a focality effect.

### Treatment effect builds up linearly

To determine the time-course of the treatment effect during the stimulation block (e.g., intra-train), we quantified the effect of each single stimulation train (from train 1 to 20) (Fig. 5d). Coherence between sensorimotor contacts was calculated in the 20 consecutive intra-train intervals during the two treatment types (8-s for iTBS; 10-s for i8Hz). The analysis window was set similar for both treatments, from +0.1-6.1 seconds after each train offset as the effects were stronger within this time interval. Figure 5d shows increased beta coherence values as a function of increasing intratrain trial for a sample electrode pair. Intra-train beta coherence values increased more strongly during iTBS compared to i8Hz treatment. Beta coherence peaks during iTBS increased steadily from 0.22 (train 1) to 0.67 (train 20) while a weaker effect occurred during i8Hz (from 0.15 in train 1 to 0.36 in train 20). We quantified the effect by fitting a linear regression line to the 20 coherence peak values and calculating the slope of the fitted function. Steeper slopes indicate larger increases in coherence values (slope iTBS = 0.0243 vs. slope i8Hz = 0.011). Across the population data (4 subjects), slopes derived from within iTBS intra-trains were steeper compared to those from i8Hz (Wilkinson signed-rank, p = 0.011) (Fig. 5e). Only contacts within the sensorimotor cortex in subjects that showed significant treatment effects were included in this analysis (subjects 1, 2, 4, and 5, 41 contacts).

A previous study [21,48] found treatment effects on CCEPs amplitude (pre vs. post). We tested this hypothesis by applying single pulses at ~1 Hz (300-ms jitter, ~200 pulses) before and after the resting periods preceding and following treatment but no treatment effects were found on the P1 amplitude of CCEPs across the population data (Wilkinson signed-rank, both iTBS and i8Hz, p > 0.2).

## Discussion

TMS is a widely used non-invasive method that stimulates large populations of neurons in masse, yet the underlying physiological mechanisms of TMS-induced synaptic plasticity are not well understood. Here, we characterized the physiological effects of direct cortical iTBS, a novel rTMS patterned stimulation applied intracranially in the human sensorimotor cortex. We found that iTBS enhanced beta coherence within sensorimotor networks. This effect was stimulation-pattern specific, as stronger effects were found using iTBS while traditional alpha burst stimulation did not increase beta coherence. This effect was limited to local, functionally connected somatosensory and motor cortical regions that share similar spectral profiles (beta oscillations) and it lasted 3 minutes after treatment offset. Our results provide a starting point for clinicians and researchers to design more optimal stimulation protocols aiming at inducing neuroplasticity in cortical networks using direct intracranial electrical stimulation.

The stimulation parameters used in this study were informed by rTMS studies aimed at inducing neuroplasticity effects [17–19,49]. The implementation of iTBS into rTMS paradigms has shortened rTMS treatment duration (40-min with conventional 10 Hz vs. 15-min with iTBS) while achieving similar or greater treatment effects [18,50]. Invasive approaches using epidural, cortical and deep brain stimulation are increasingly used to treat movement/neuropsychiatric disorders and epilepsy [22,51,52] and investigated to assist motor recovery following strokes and traumatic brain injuries [1,8,53]. Although the exact mechanisms underlying treatment effects using these invasive stimulation approaches are still largely unknown, neuroplastic changes are thought to play a significant role [13,14,54,55]. The goal of our study is to determine the most ideal stimulation parameters to induce cortical neuroplasticity in humans.

Prior studies translating rTMS paradigms to intraparenchymal stimulation showed varying or weak results [21,23], or did not report neurophysiological changes [36]. For example, Keller et al. (2018) stimulated several different brain regions across patients and found mixed excitation at some treatment locations and inhibition at others. The authors found that intermittent alpha frequency stimulation of the DLPFC produced excitation in 2 subjects and suppression in the other 2, while stimulation of temporal (1 subject) and motor (3 subjects) cortical regions produced suppression. In our study we limited the treatment (and read-out) sites to one well defined functional network (sensorimotor), and measured changes in excitability after trains of theta bursts (iTBS). Our results indicate that iTBS can enhance neuronal plasticity more effectively compared to alpha frequency stimulation (i8Hz). An additional interesting result from our study was that, unlike Keller et al., we did not find significant effects on CCEPs but on spontaneous beta coherence during resting state, indicating that this measure may be more sensitive to treatment-related changes. Another difference was that we did not observe significant effects on CCEPs in the motor cortex after trains of i8Hz stimulation, however, our stimulation currents were applied at lower amplitudes (80% of the active motor threshold vs. 100%) and for shorter periods of time.

A key result from our work is that iTBS treatment increased coherence specifically within the sensorimotor system and at beta (12-30 Hz) frequencies. Importantly, areas within the sensorimotor cortices already exhibited prominent coherence peaks in the beta band prior to treatment (Fig. 2b and 2c), suggesting that iTBS enhanced connectivity of already pre-existent rhythms. Prior studies showed that the frequency of stimulation is an important factor in driving specific clinical outcomes in Parkinson’s patients [10,36,56,57], in epilepsy patients [15,51], in modulating focal excitability in macaque brains [48], and in prolonging the duration of the effects (acute vs. plasticity) [52,58,59]. The focal spread of our treatment effects is supported by invasive studies that showed the majority of modulated regions being anatomically and functionally closer to the stimulation site [21,34]. Regarding the focal vs. network specificity of the effect, electrode contacts that were within sensorimotor areas and near the treatment site showed the strongest iTBS effects. However, contacts that were more distant from each other yet within sensorimotor areas (medial vs. lateral portions of the motor cortex) also showed increased beta coherence compared to other equally distant contacts that were outside sensorimotor areas (e.g., insula). This result suggests a network specific effect rather than just a focality effect.

Since we characterized the physiological effects of cortical iTBS in the human sensorimotor cortex the obvious question is about the generalizability of the results. A recent DBS study in PD patients applied subcortical iTBS to the globus pallidus internus (GPi) and the subthalamic nucleus (STN) while recording from the DLPFC [23]. They found that GPi stimulation (but not STN) modulated theta power in DLPFC, suggesting that iTBS can be effectively applied to other brain areas and cause frequency-specific effects (see also [60]). In iTBS, trains of pulses are delivered at 5 Hz which raises the question as to why a resonance frequency of ~5 Hz amplifies coherence in the beta range in our results. Further research will need to confirm whether iTBS can amplify pre-existent rhythms in specific brain areas or predominantly theta rhythms [60], or whether alternative approaches using close-loop stimulation [48,61] and/or adaptive DBS [10] will be more efficacious.

Our effects induced by cortical iTBS treatment outlasted the period of stimulation, as beta coherence remained elevated for ~3 minutes after treatment offset. Studies in mice hippocampus have convincingly shown long-lasting synaptic potentiation (LTP) effects outlasting iTBS for several hours and even days. However, iTBS treatments in animal models are delivered multiple times during hours, repeated over several days and at high amplitude currents. Recent human rTMS studies suggested that a single iTBS therapy session is insufficient and that multiple sessions might provide benefit [26,27]. Our findings indicate that multiple iTBS treatments do not result in linear increases in beta coherence in line with a recent study [25], suggesting more complex dynamics that likely are subject and activity-dependent [62]. Further cortical iTBS studies might confirm the number of sessions required to induce longer lasting effects in the human cortex. Analysis of activity within a single treatment session (intra-train stimulation intervals) might clarify the time-course of the treatment effects [34]. Our results show that iTBS mediated beta coherence effects scale more or less linearly with consecutive trains of stimulation (Fig. 5d). One advantage of the invasive cortical stimulation used here compared to rTMS is that the electrodes and the stimulator could be implanted and thus treatment could be applied continuously.

Our iTBS treatments were applied during resting-state while patients were quietly relaxed, and no behavioral output was measured. Future studies might investigate whether intracranial sensorimotor cortex iTBS influences resting motor threshold (RMT) or MEP amplitude, and whether it can improve the execution and learning of specific motor tasks. Two DBS human studies applied TBS in the entorhinal white matter and found improvements in memory for portraits, while stimulation of the adjacent entorhinal grey matter of the subiculum did not improve memory [28,63]. Most of our patients (6/7) were implanted with sEEG arrays, and stimulation was applied through 2 contacts in a bipolar fashion spanning a total distance of 10-mm (Fig. 2a), which likely stimulated passing-by white matter tracts, contributing to the treatment effect [15]. In fact, some of the contacts used for treatment were in the white-grey matter junction of the sensorimotor cortex. This might explain the null effect observed in patient 6, who was implanted with ECoG grids, as subdural stimulation spreading along the cortical convexity might less effectively excite white matter tracts compared to sEEG stimulation. This explanation cannot account for subjects 3 and 7 who were implanted with sEEG and showed no significant treatment effects. We did not find any systematic difference regarding treatment locations, proximity to white matter tissue, or spectral frequency profiles in these subjects. Current rTMS treatments are efficacious in approximately 30% of patients which is in line with our results. Further work comparing DBS programming strategies in bigger data sets [64] and electrical field modelling [65,66] is needed to improve the interpretability of the results and reduce the wide interindividual differences regarding the ideal stimulation site relative to the cortex.

A potential pitfall of this study is that it was conducted in patients with epilepsy. Future studies will have to establish if the results generalize to other patient populations. To increase potential generalizability and due to safety precautions, we only included patients with their epileptogenic focus outside of the sensorimotor system and at least 1 gyrus removed from the treatment site (Table 1).

## Conclusions

The therapeutic potential of rTMS patterned stimulation paradigms using iTBS has revolutionized the field of electronic medicine. Improved treatment efficacy of iTBS compared to conventional rTMS was shown in a variety of neurological disorders including PD, epilepsy, depression, OCD and PTSD. Despite these promising results, a clear understanding of the underlying neurophysiological mechanisms in humans is lagging that observed in animal models where iTBS induces robust long-lasting effects on glutamatergic synapses. We bridge this gap by applying iTBS directly into the cortex of patients implanted with depth electrodes for epilepsy monitoring. We found that cortical iTBS induces stronger effects compared to conventional intermittent alpha burst stimulation, and that these effects are frequency and spatially specific. Specifically, iTBS applied to well-defined regions of the sensorimotor cortex at low amplitude currents increases pre-existent local synchrony in the beta range. In summary, iTBS can enhance neuronal plasticity more effectively compared to other treatment modalities within a single experimental session. Our results indicate that iTBS enhancement is frequency and amplitude dependent, and that it occurs within local sensorimotor networks that share similar spectro-temporal properties (beta oscillations). Our findings might help explain part of the somewhat heterogeneity in the results across studies using repetitive stimulation and strongly suggest that standard iTBS protocols (either non-invasive or invasive) consider the individuals excitability profile (e.g., pre-existent rhythms in a given cortical area).

## Abbreviations

iEEG: intracranial electroencephalogram
iTBS: Intermittent Theta Burst Stimulation
i8Hz: Intermittent Alpha Burst Stimulation
DBS: Deep Brain Stimulation
rTMS: repetitive Transcranial Magnetic Stimulation
CCEPs: Cortico-cortical evoked potentials
HFSM: High Frequency Stimulation Mapping
HFA: High Frequency Activity
LTP/LTD: Long-term Potentiation/Suppression

## Author Contributions

**Jose L. Herrero**: Conceptualization, Methodology, Investigation, Analysis, Writing, Supervision. Visualization. **Alexander Smith:** Data curation, Visualization, Writing. **Akash Mishra:** Data curation. **Noah Markowitz**: Data curation. Visualization. **Ashesh D. Mehta**: Conceptualization, Supervision, Funding acquisition. Writing. **Stephan Bickel:** Conceptualization, Methodology, Investigation, Supervision, Writing.

## Acknowledgements

We are indebted to all patients who volunteered their time to participate in our study. We also thank the staff of the Epilepsy Monitoring Unit at North Shore University Hospital for their support throughout the conduction of the studies. We thank Nima Mesgarani and Stavros Zanos for providing helpful feedback on the manuscript. We thank Tucker-Davis Technologies (Mark Hanus and Myles Billard) for support with the neural recording and stimulation equipment

